# Restoration of mitochondrial structure and function within *Helicobacter pylori* VacA intoxicated cells

**DOI:** 10.1101/2023.05.19.540400

**Authors:** Robin L. Holland, Kristopher D. Bosi, Ami Seeger, Steven R. Blanke

## Abstract

The *Helicobacter pylori* vacuolating cytotoxin (VacA) is an intracellular, mitochondrial-targeting exotoxin that rapidly causes mitochondrial dysfunction and fragmentation. Although VacA targeting of mitochondria has been reported to alter overall cellular metabolism, there is little known about the consequences of extended exposure to the toxin. Here, we describe studies to address this gap in knowledge, which have revealed that mitochondrial dysfunction and fragmentation are followed by a time-dependent recovery of mitochondrial structure, mitochondrial transmembrane potential, and cellular ATP levels. Cells exposed to VacA also initially demonstrated a reduction in oxidative phosphorylation, as well as increase in compensatory aerobic glycolysis. These metabolic alterations were reversed in cells with limited toxin exposure, congruent with the recovery of mitochondrial transmembrane potential and the absence of cytochrome *c* release from the mitochondria. Taken together, these results are consistent with a model that mitochondrial structure and function are restored in VacA-intoxicated cells.

## INTRODUCTION

Mitochondria regulate a diverse array of cellular functions, including the maintenance of metabolic homeostasis (1), initiation of programmed cell death (1), and intracellular innate immunity (2). The importance of mitochondria for cellular health in humans is exemplified by the association between mitochondrial dysfunction and diverse human disorders and diseases, including Parkinson’s disease (3), Leber’s hereditary optic neuropathy (4), and tumorigenesis (5). Among the known etiologies associated with mitochondrial dysfunction, pathogenic microbes have increasingly been recognized to impact mitochondrial health and function (6), as an integral component of their virulence strategies (2, 7). For example, *Neisseria gonorrhoeae* (8), *Neisseria meningitidis* (9), and *Escherichia coli* (10), interfere with mitochondrial membrane integrity in a manner that results in either activation (9, 10) or inhibition (8) of mitochondrial-dependent cell death pathways. In addition, other pathogenic agents, such as Influenza A (11), Hepatitis C (12), and *Vibrio cholerae* (13), usurp mitochondrial function as a means to evade innate immunity.

One of the first microbes identified to target mitochondria in eukaryotic cells was *Helicobacter pylori* (*Hp*), which is an important and broadly distributed gastric pathogen of humans. Chronic infection with *Hp* is the single most important risk factor associated with the onset of gastric cancer. Among the virulence factors that promote colonization, persistence, and changes to the gastric mucosa-associated with malignancy, the vacuolating cytotoxin (VacA) is the only known intracellular-acting exotoxin secreted by *Hp*. The importance of VacA as a risk factor for gastric disease is supported by human epidemiological evidence (14–17). In addition, animal studies have demonstrated the importance of VacA for the colonization of *Hp* within the gastric glands of the infected stomach (18, 19). *In vitro* cell culture studies have revealed alterations that occur in gastric epithelial and immune cells exposed to VacA, including vacuole biogenesis (20, 21), mitochondrial dysfunction (22–26), the induction of autophagy (27–29), and cell death (26, 30–32).

Although the exact role of VacA as an important determinant of *Hp* virulence during chronic infection remains poorly understood, there is growing evidence that VacA alters overall metabolism within intoxicated cells. Several studies have indicated that following uptake into cultured cells, VacA is localized to mitochondria (22, 26, 33, 34), and more specifically, to the inner mitochondrial membrane (25, 33, 34), which becomes depolarized in a manner dependent upon the toxin’s membrane channel-forming activity (22, 23). Dissipation of mitochondrial transmembrane potential (ΔΨ_m_) leads to the collapse of proton motive force, which is critical for maintenance of cellular ATP levels (23, 24, 35). In addition, mitochondrial dynamics are disrupted in cells exposed to VacA, leading to excess fragmentation of mitochondrial structure (35). Alterations in mitochondrial structure and function are sensed globally within VacA intoxicated cells, as indicated by toxin-dependent inhibition of the mammalian target of rapamycin complex-1 (mTORC1), prompting a shift in cellular metabolic priorities towards catabolism through, in part, the induction of autophagy (36). Despite advances in our understanding of VacA-dependent mitochondrial modulation, the extended consequences of toxin-mediated mitochondrial dysfunction, including host responses to intracellular VacA action, are poorly understood.

Here, we report that VacA-dependent mitochondrial damage is limited within host cells and, under experimental conditions where toxin exposure is restricted, both organelle structural and function were restored in a time-dependent manner. Collectively, results from these studies suggest that host cells respond to *Hp* modulation of host cell metabolism as a consequence of intracellular VacA activity.

## RESULTS

### Restoration of filamentous mitochondrial structure within VacA intoxicated cells

Fragmentation of mitochondria occurs in response to cellular stress (37–40), but is also part of normal cell physiology (41, 42). Exposure to VacA has previously been linked to the rapid fragmentation of filamentous mitochondria as a result of mitochondrial dynamics modulation (35). However, the effects of extended cellular exposure to VacA on mitochondrial structure, as well as mitochondrial function and overall cellular viability is incompletely understood. To better understand the consequences of extended exposure to VacA-mediated targeting of mitochondria, we first monitored mitochondrial structure in cells exposed continuously to toxin for up to 24 h. AZ-521 duodenal-derived epithelial cells, which have been used extensively to study VacA-mediated alterations in cell culture (24, 35, 43), were exposed to VacA that had been purified from culture filtrates from *Hp* 60190, a clinical isolate that has been widely used to study VacA biology (31, 36, 44, 45). Monolayers of AZ-521 cells were preincubated at 4 °C, which is not permissive for uptake of membrane-bound surface components into cells. After 30 min, monolayers were further incubated at 4 °C with VacA (250 nM) to promote binding of toxin to the cell surface in the absence of cellular uptake. After an additional 30 min, the monolayers were shifted to 37 °C, which is permissive for toxin internalization, and will hereafter be referred to as “continuous exposure to prebound toxin”. After 1, 4, and 24 h, the monolayers were examined for mitochondrial structure using fluorescence microscopy. These experiments revealed that mitochondria were both visibly and quantitatively more fragmented in monolayers that had been exposed for 1 h to VacA, than in monolayers exposed to the vehicle control (PBS pH 7.4) (Fig. 1), as previously reported (35). Likewise, after 4 h, mitochondria remained more highly fragmented in cells continuously exposed to VacA, relative to vehicle-treated cells. However, after 24 h, long filamentous mitochondrial structures, and fewer fragmented mitochondria, were readily evident in monolayers that had been exposed continuously to VacA. The partial restoration of mitochondrial structure in cells continuously exposed to toxin suggested that VacA intoxicated cells were able to limit the effects of continuous toxin exposure on mitochondrial structure.

**Figure 1.**
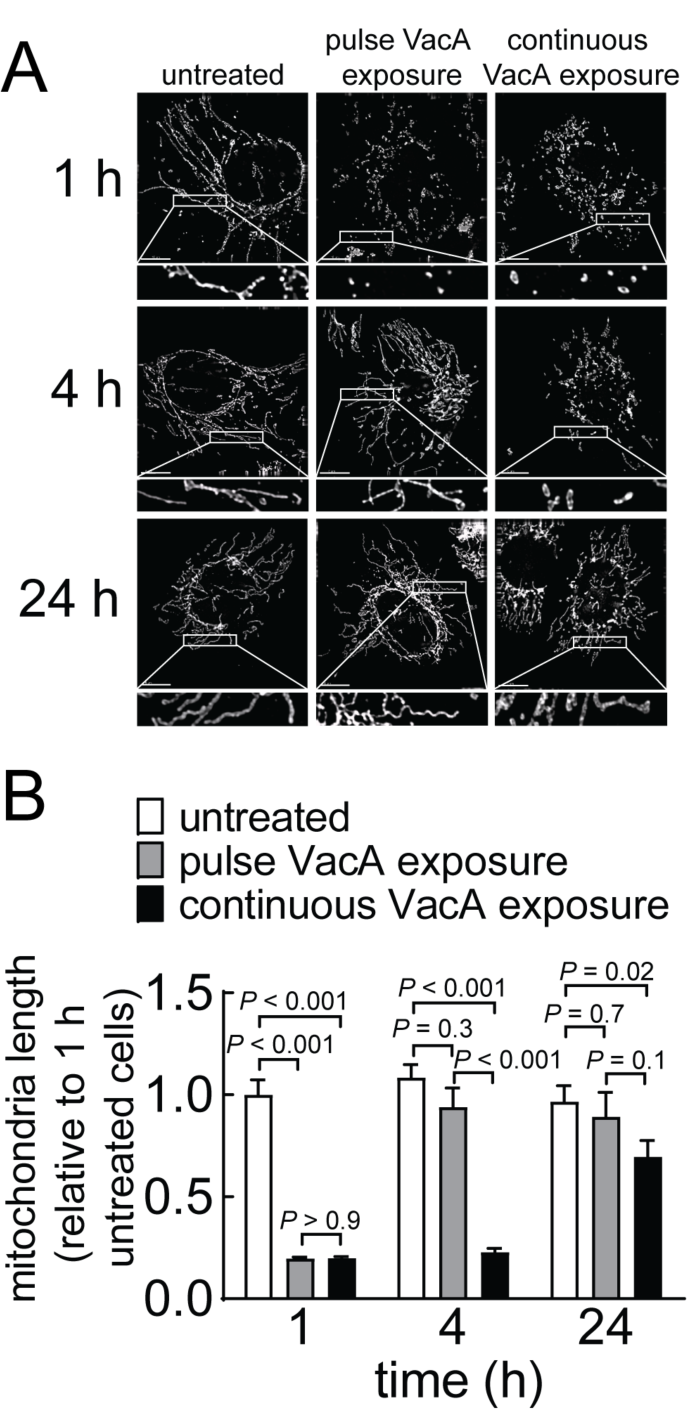
Mitochondria undergo transient fragmentation within cells exposed to VacA. AZ-521 cells were incubated in the absence or presence of VacA (250 nM) under “cold” pulse exposure conditions (4 ⁰C for 30 min), washed with PBS (pH 7.2), and incubated at 37 °C under 5% CO_2_ in fresh cell culture medium in the absence of VacA. Alternatively, AZ-521 cells were preloaded by incubating at 4 °C in the absence or presence of VacA (250 nM). After 30 min, cells were further incubated continuously at 37 °C. After 1, 4, or 24 h for both “cold” pulse exposure and continuous exposure conditions, monolayer cells were fixed, permeabilized, and changes in mitochondrial structures were visualized by incubating the monolayers with anti-TOM20 antibodies, followed by incubation with fluorescently labeled secondary antibodies, and imaged by fluorescence microscopy. (B) Mitochondrial length was measured in cells that had been administered VacA, relative to mitochondria within untreated cells. Scale bar represents 10 µm. Images in (A) are representative of those collected over three independent experiments (n=3). Results presented in (B) were derived from data combined from three independent experiments (n=3). Error bars represent standard error of the mean. Statistical significance was determined using one-way ANOVA with alpha of 0.001 (α=0.001), with the Tukey’s correction applied for multiple comparisons against the untreated control within each timepoint.

### Filamentous mitochondrial structure is restored more quickly in cells with limited exposure to VacA

The results described immediately above suggested the possibility that cells were able to limit toxin-dependent mitochondrial structural alterations resulting in partial restoration of filamentous mitochondria. If this is true, we predicted that restoration of filamentous mitochondrial structure would occur more quickly in cells with limited exposure to toxin, which hereafter will be interchangeably referred to as “pulse exposure”. Monolayers of prechilled cells were incubated at 4 °C in the absence or presence of VacA (250 nM). After 30 min, the monolayers were washed with ice-cold PBS pH 7.4, to remove unbound-VacA, and then further incubated at 37 °C with complete medium in the absence of toxin. Similar to the results obtained from studies conducted under conditions of continuous exposure to prebound toxin (Fig. 1), fluorescence microscopy revealed that cells with limited exposure to VacA for 1 h possessed visibly and quantitively more fragmented mitochondria than monolayers that had been exposed for 1 h to vehicle (Fig. 1). However, in contrast to monolayers continuously exposed to VacA, mitochondria were predominantly filamentous at 4 h within cells after limited exposure to the toxin. After 24 h, cells that had been transiently exposed to VacA possessed filamentous mitochondria, similar to cells within monolayers that had been incubated with the vehicle control, further supporting the idea that cells exposed to VacA are able to not only limit, but in fact counteract the effects of toxin action on mitochondrial structure. These results obtained in AZ-521 cells were similar to those obtained in adenocarcinoma gastric cells (AGS) (Fig. S1), which have been extensively used as an *in vitro* culture model for studying VacA biology (30, 46–49). Overall, these results suggest that cells exposed to VacA offset the effects of toxin action on mitochondrial structure.

### Restoration of mitochondrial transmembrane potential (ΔΨ_m_) and ATP levels in cells with continuous or limited exposure to VacA

Having demonstrated, for the first time, that VacA-dependent alterations in mitochondrial structure are reversible, we next evaluated whether toxin-mediated mitochondrial dysfunction might also be reversible within intoxicated cells. Experiments to evaluate time-dependent changes in mitochondrial transmembrane potential (ΔΨ_m_), which is required for oxidative phosphorylation and ATP generation, revealed significantly lower ΔΨ_m_ levels in cells that had been exposed to VacA continuously for 1, 4, and 24 h, with the levels of ΔΨ_m_ remaining constant between 4 and 24 h (Fig. 2A).

**Figure 2.**
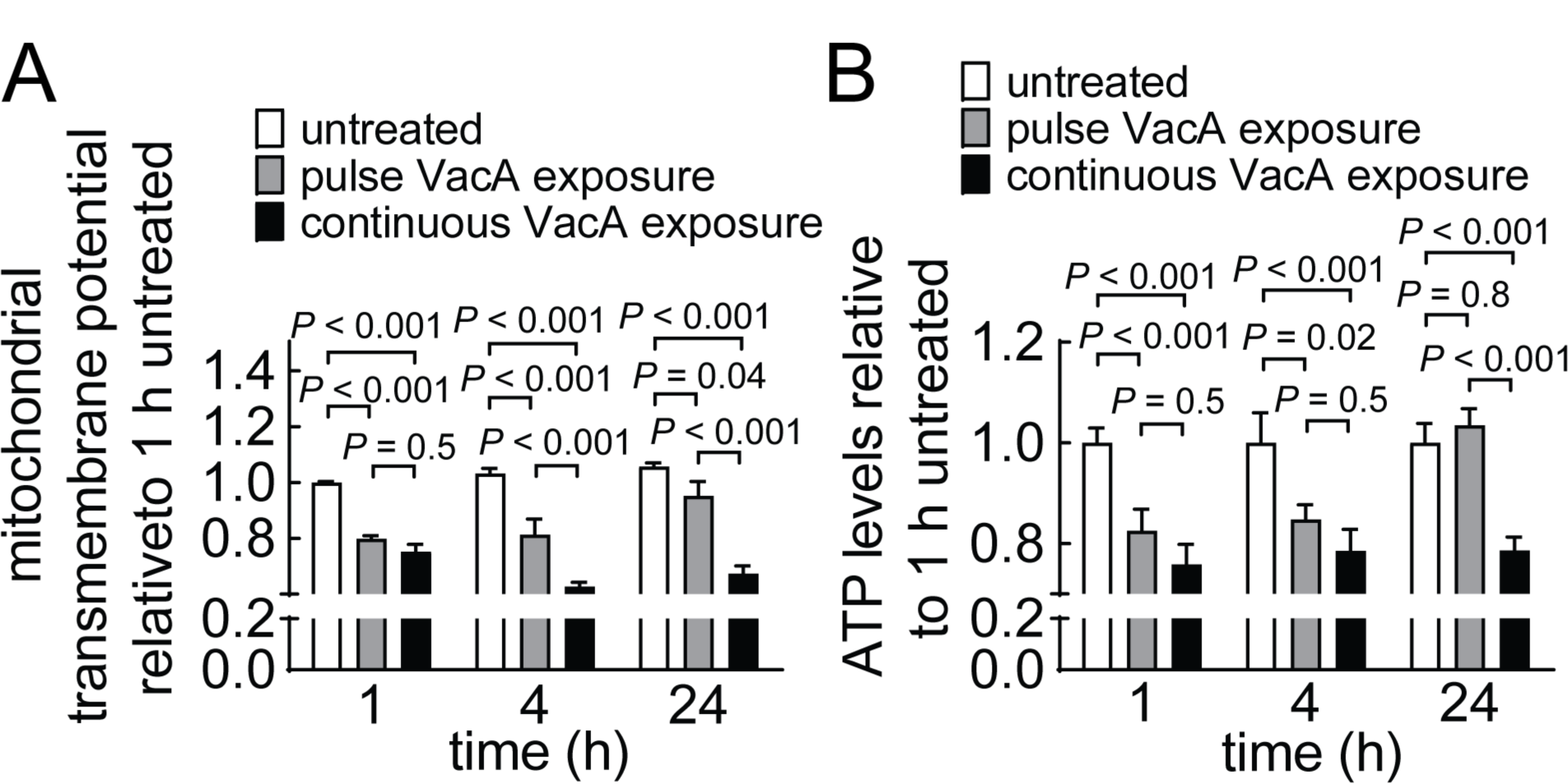
VacA-mediated changes in mitochondrial function are dynamic. AZ-521 cells were incubated in the absence or presence of VacA (250 nM) under “cold” pulse exposure conditions or continuous exposure conditions. After 1, 4, or 24 h, cells were collected for analyses. (A) Mitochondrial transmembrane potential was measured using flow cytometry of cells stained with TMRE (10 nM, 30 min), with 10,000 events collected for each individual treatment. (B) ATP levels were measured using the MitoTox kit, as described under Materials and Methods, with normalization to total protein. Data were combined from three independent experiments, each performed in triplicate. Error bars represent standard error of the median (A) or mean (B). Statistical significance was determined by one-way ANOVA with alpha of 0.001 (α=0.001), using the Tukey’s multiple comparisons test against the untreated control at each timepoint.

After 1 h, cells with pulse exposure to VacA, showed similar ΔΨ_m_ dissipation as cells exposed continuously to toxin. However, after 4 and 24 h, significantly higher ΔΨ_m_ was detected in monolayers with limited exposure to VacA relative to continuous exposure to the toxin. Similar trends were observed in AGS cells (Fig. S1). From these results, we conclude that VacA-dependent depolarization of the mitochondrial inner membrane is reversible, resulting in restoration of mitochondrial transmembrane potential, which is required for energy production during oxidative phosphorylation.

In further support of this idea, experiments to evaluate time-dependent changes in cellular energy, revealed lower levels of cellular ATP in monolayers that had been either pulse or continuously exposed for 1 or 4 h to VacA relative to treatment with vehicle (Fig. 2B). After 24 h, cells continuously exposed to VacA maintained decreased levels of ATP, similar to that measured at 1 and 4 h, indicating that cellular ATP levels remained constant. In contrast, cellular ATP levels were higher in cells with limited exposure to VacA for 24 h, similar to vehicle-treated (untreated) cells. These results further support the idea that, within cells exposed to VacA, full reestablishment of ΔΨ_m_ at the mitochondrial inner membrane resulted in restoration of cellular energy.

Under conditions of normal metabolic function, cells efficiently generate ATP through oxidative phosphorylation, which requires highly functioning mitochondria (50). Experiments were performed to address the effects of VacA-mediated mitochondrial dysfunction on oxidative phosphorylation through the measurement of oxygen consumption (as a surrogate for oxidative phosphorylation). These studies revealed significantly lower levels of oxygen consumption by monolayers incubated continuously with toxin for 4 and 24 h in the presence than the absence of VacA (Fig. 3A). In contrast, significant differences in oxygen consumption were not detected after 1, 4, or 24 h in monolayers with limited exposure to VacA. These results suggest that cells function to limit toxin-mediated reduction in oxidative phosphorylation. In addition, restoration of mitochondrial function occurs sooner in cells with limited exposure to VacA relative to cells continuously exposed to the toxin. These findings are congruent with the idea that overall mitochondrial damage is dictated by flux of mitochondrial-associated VacA, which represents the balance of mitochondrial exposure to toxin versus cellular restoration of organelle function.

**Figure 3.**
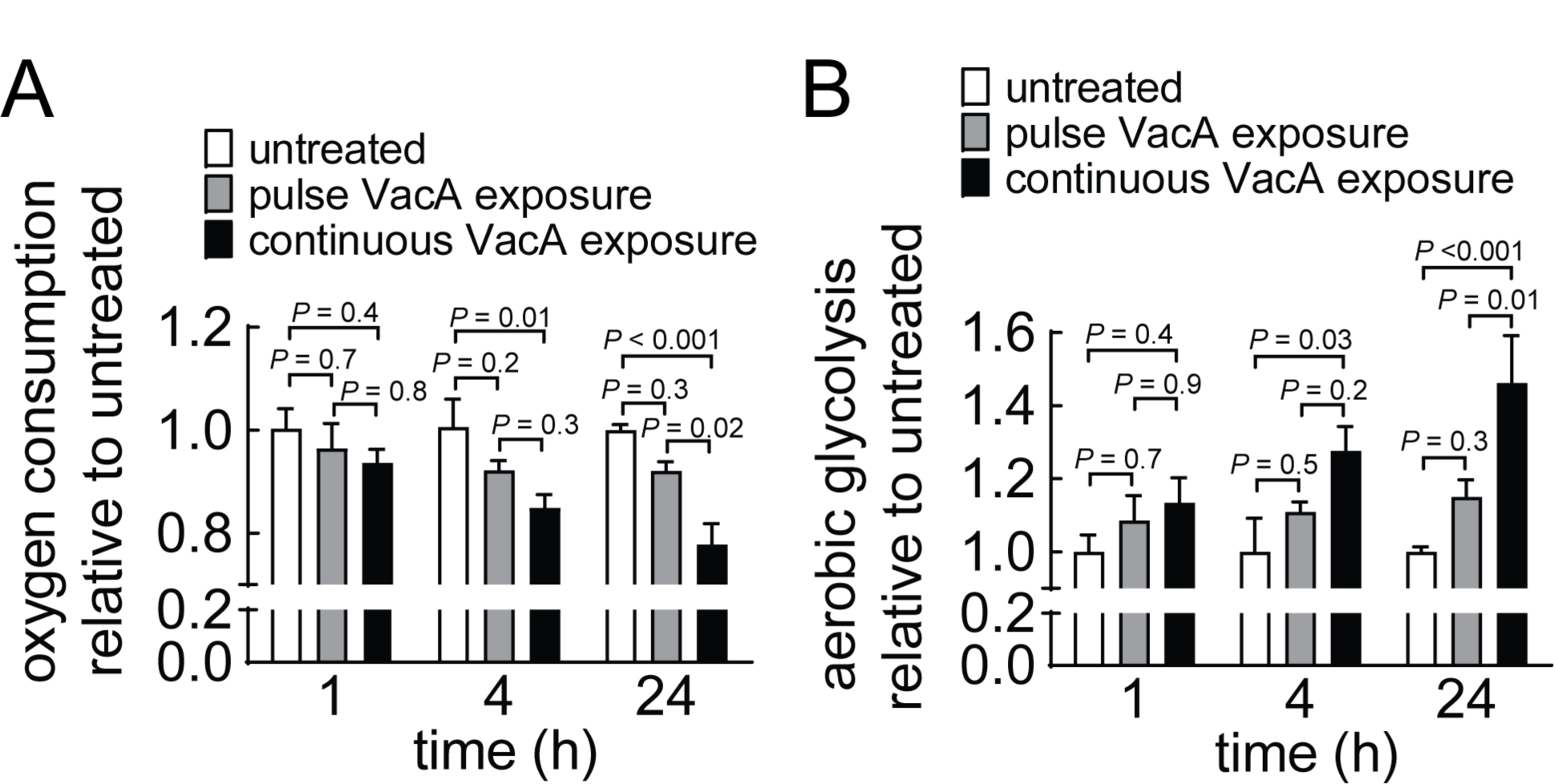
VacA-mediated changes in mitochondrial function are associated with a compensatory shift from oxidative phosphorylation to aerobic glycolysis. AZ-521 cells were incubated in the absence or presence of VacA (250 nM) under “cold” pulse exposure conditions or continuous exposure conditions. After 1, 4, or 24 h, cells were collected for analyses. Oxygen levels (A) and aerobic glycolysis (B) were measured using Oxygen Consumption/Glycolysis Dual Assay Kit or the MitoXpress kit, as described under Materials and Methods. Data were combined from three independent experiments each performed in triplicate. Error bars represent standard error of the mean. Statistical significance was determined by one-way ANOVA with alpha of 0.001 (α=0.001), using Tukey’s correction for multiple comparisons against the untreated control at each timepoint.

To further validate the metabolic changes resulting from VacA-dependent mitochondrial dysfunction, we examined cells that had been exposed to VacA for an increase in aerobic glycolysis, which can partially compensate for reduced energy production with generation of ATP, albeit at substantially lower levels than those resulting from oxidative phosphorylation (51). To evaluate whether aerobic glycolysis is upregulated in cells exposed to VacA, we examined VacA-dependent cellular production and secretion of lactic acid, which is a metabolic byproduct of aerobic glycolysis (51–53). These studies revealed significantly higher levels of lactic acid in the medium collected from monolayers incubated continuously for 4 and 24 h in the presence than absence of VacA (Fig. 3B). In contrast, significant increases in lactic acid levels were not detected after 1, 4, or 24 h in monolayers transiently exposed to VacA. These studies further corroborate a model that, in response to mitochondrial dysfunction within cells intoxicated with VacA, shifts in metabolism occur that are consistent with impaired energy production. Moreover, these results are also consistent with the idea that cells can limit and/or reverse the extent to which energy production is impaired.

### Mitochondrial outer membrane integrity is not compromised in cells with limited exposure to VacA

Previous studies indicated that continuous exposure to VacA results in an increase in mitochondrial outer membrane permeability (MOMP), which releases cytochrome *c* from the intermembrane space of mitochondria into the cytosol, thereby committing the cell to mitochondrial-associated programmed cell death (22, 23). To further evaluate the effects of the relationship between mitochondrial dysfunction and MOMP in cells with limited exposure to VacA, we examined cytochrome *c* release using immunofluorescence microscopy in cells that had been exposed to the toxin in either a continuous or limited manner. These studies showed that cells that had been pulse exposed to VacA for 24 h did not undergo MOMP, as cytosolic cytochrome *c* was not detected within any of the cells analyzed (Fig. 4). There was also no evidence of MOMP-mediated release of cytochrome *c* into the cytosol in cells that had been continuously exposed to VacA for 1 or 4 h. However, after 24 h, cytochrome *c* release into the cytosol was detected in only 20.5 (± 11.4%) of the cells continuously exposed to VacA (Fig. 4). These results suggest that restoration of mitochondrial function, as indicated by an increase in oxidative phosphorylation and cellular ATP, as well as a decrease in aerobic glycolysis, has a positive association with survival of cells that have been intoxicated with VacA.

**Figure 4.**
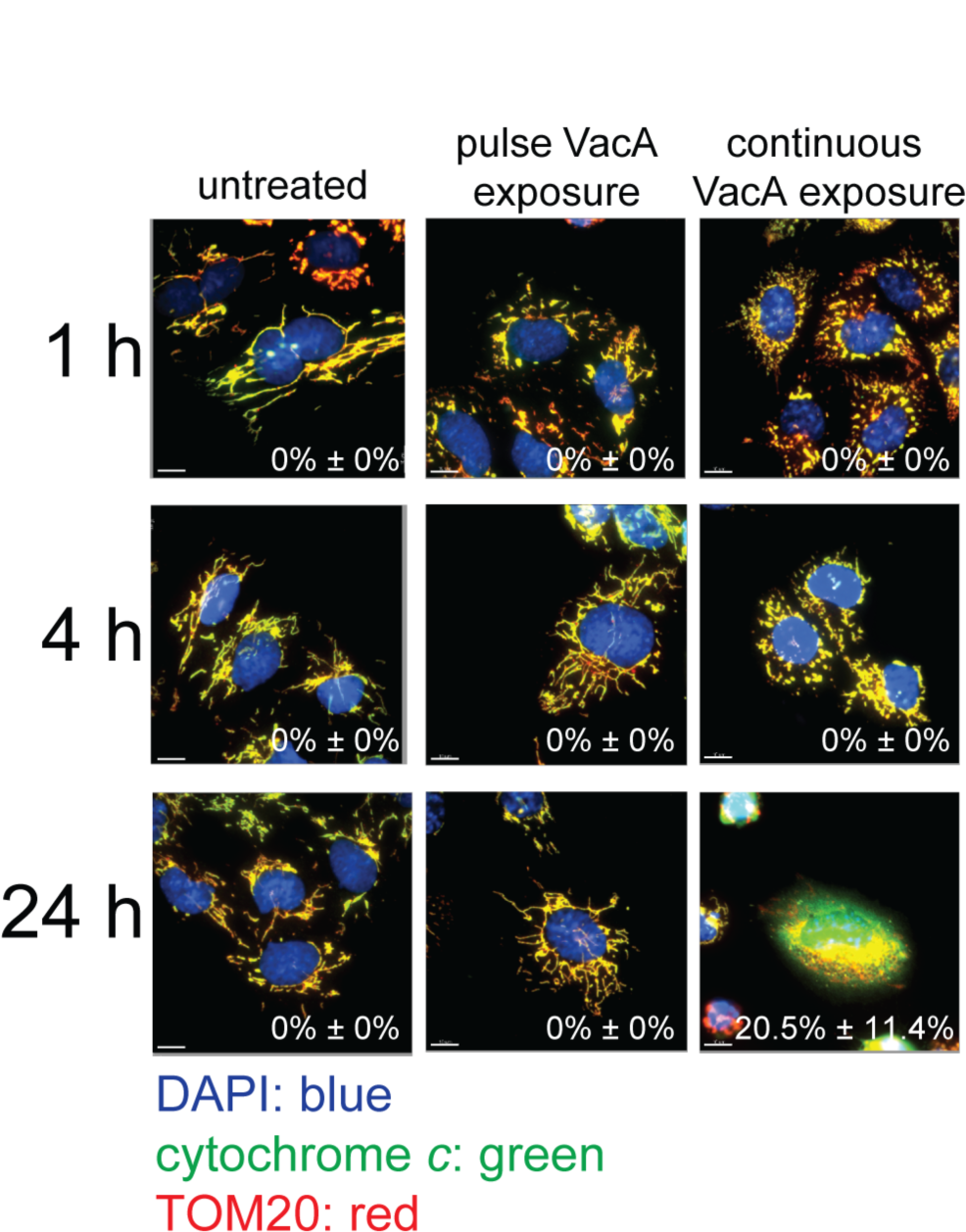
VacA-mediated changes in mitochondrial function are dynamic. AZ-521 cells were incubated in the absence or presence of VacA (250 nM) under “cold” pulse exposure conditions or continuous exposure conditions. After 1, 4, or 24 h, cells were collected for analyses. Cells were stained with anti-TOM20 and anti-cytochrome *c* antibodies, followed by incubation with fluorescently labeled secondary antibodies, and imaged by fluorescence microscopy. Scale bars represent 10 µm. The values notated on the bottom right corner of each image represent percentage of cells displaying cytochrome *c* release, assessed by fluorescence microscopy (mean ± SD). Images are representative of three independent experiments each performed in triplicate. The percentages were combined from three independent experiments each performed in triplicate. Statistical significance was determined by one-way ANOVA with alpha of 0.001 (α=0.001), using Tukey’s correction for multiple comparisons against the untreated control within each timepoint.

## DISCUSSION

While VacA targeting of mitochondria has been previously reported to alter overall cellular metabolism, little is known about the consequences of extended exposure to the toxin. Consistent with previous reports (23, 24), results from experiments described here validated that continuous exposure of cultured epithelial cells to VacA for 1 h induced mitochondrial dysfunction, as indicated by a significant loss in mitochondrial transmembrane potential (DY_m_) (Fig. 2A) and reduction in cellular ATP (Fig. 2B). However, further significant dissipation of DY_m_ and reduction of cellular ATP did not occur in monolayers continuously exposed to VacA for 4 and 24 h, suggesting that VacA-dependent mitochondrial damage within intoxicated cells had been limited. One possible explanation for the significant but limited mitochondrial dysfunction in response to VacA is that cells possess a mechanism for responding and restricting VacA-mediated effects at this organelle, even as cells continue to be exposed to toxin. If this is true, we predicted that exposing cells to a “single pulse” of VacA would result not only in restricting VacA-dependent dysfunction but would also lead ultimately to restoration of mitochondrial function. Although the dissipation of DY_m_ and reduction of cellular ATP occurred to approximately the same extent in cells that had been either continuously-or pulse-exposed to VacA for 1 h, monolayers pulse-exposed to VacA showed significantly higher levels after 4 and 24 h, indicating that mitochondrial function had been restored in cells that had only limited exposure to the toxin (Fig. 2A and B).

The mechanism by which cells orchestrate restoration of mitochondrial function within VacA intoxicated cells remains to be established. Previous studies have reported that following uptake of VacA into sensitive cells, there is an increase in mitochondrial-associated toxin (23, 26, 33, 34). Localization of VacA to the mitochondrial inner membrane has been previously reported, which is consistent with a current model that VacA forms ion-conducting channels responsible for dissipation of the inner mitochondrial transmembrane potential (23), resulting in the collapse of proton motive force required for ATP production (24). On the other hand, in one recent report, the authors did not observe substantial accumulation of mitochondrial-associated VacA over time (54). While the significance of this observation is yet not known, it is tempting to speculate that upon initial VacA-dependent mitochondrial dysfunction, cells might invoke a mechanism to limit and restore mitochondrial function by restricting and/or actively removing mitochondrial-associated VacA. Ongoing work in our laboratory is focused on identifying the mechanism underlying the capacity of cells to efficiently counteract the mitochondrial damaging action of the toxin.

In addition to limiting and reversing VacA-dependent mitochondrial dysfunction, our work also revealed that filamentous mitochondria are restored within intoxicated cells. Mitochondrial fragmentation has been recognized as a consequence of VacA intoxication for a number of years (35) and is considered generally as a response to mitochondrial stress (37–40). Our studies revealed significant fragmentation of mitochondria in cells that had been either continuously-or pulse-exposed to VacA for 1 h (Fig. 1). Notably, after 4 h, cells that were pulse-exposed to VacA had recovered their filamentous phenotype, while cells that had been continuously exposed to toxin remained predominantly fragmented. However, after 24 h of continuous exposure to toxin, the cells exhibited almost exclusively filamentous mitochondria. Our data indicate that mitochondrial structure and function are both restored within VacA intoxicated cells, albeit to differing degrees. These results also suggest that recovery of mitochondrial structure precedes restoration of mitochondrial function, although the relationship between these two phenomena within VacA intoxicated cells remain poorly understood.

The exact role of VacA-mediated mitochondrial targeting during *Hp* gastric infection remains incompletely understood. Administration of crude VacA-containing *Hp* extracts into the stomachs of mice by oral gavage was earlier reported to cause readily evident histological damage, including wide-spread cell death within the gastric mucosa (55–59). Multiple studies have indicated loss of cell viability in gastric cells exposed to VacA in both a concentration- and time-dependent manner (22, 30–32, 35, 60–63). Although several mechanisms of cell death have been described, the capacity of VacA to induce mitochondrial outer membrane permeability (MOMP), as indicated by the released of cytochrome *c* from the mitochondrial intermembrane space into the cytosol, suggests that cell death can occur by a mitochondrial-dependent mechanism of programmed cell death (64). Much of the previous *in vitro* cell culture work with VacA has been conducted in the presence of the weak base NH_4_Cl, which recent work suggests may be primarily responsible for cell death that historically has been associated with the toxin (54). In fact, the continuous gastric infusion of purified VacA into the stomachs of mice for extended periods of time (up to 30 days), did not result in physiological damage to the gastric mucosa, or evidence of increased cell death (65). Although high concentrations of VacA may cause low levels of cell death *in vitro*, the primary role of VacA during infection is likely not the killing of host cells. Rather, it seems that VacA functions primarily in epithelial cells to modulate cellular metabolism by causing mitochondrial dysfunction. The balance between mitochondrial dysfunction and restoration could provide a means for the toxin to impede host cell metabolism without ultimately killing host cells.

In summary, we report here that not only do mammalian cells mostly survive the effects of VacA-dependent mitochondrial dysfunction, but that VacA-intoxicated cells also demonstrate the capacity to limit and restore mitochondrial structure and function. Ongoing work in our laboratory is focused on identifying mechanisms underlying the capacity of cells to efficiently counteract the mitochondrial damaging action of this toxin.

## MATERIALS AND METHODS

### Bacterial strains

*Helicobacter pylori (Hp)* 60190 (*cag* PAI^+^, *vacA* s1/m1; ATCC 49503) was cultured in bisulfite/sulfite-free *Brucella* (BSFB) broth (4 mL, 0.5% NaCl, 0.2% β-cyclodextrin, 0.1% dextrose (all from Sigma-Aldrich, St. Louis, MO), and, 1% peptone, 1% tryptone, and 0.2% yeast extract (all from BD Bacto, NJ), overlaid on 2% agar plates made of Ham’s F-12 medium (Sigma-Aldrich) supplemented with 5% fetal bovine serum (FBS) (Sigma-Aldrich)), and incubated in a microaerophilic environment (10% O_2_, 5% CO_2_).

### Mammalian cells

AZ-521 human duodenal adenocarcinoma derived cells were obtained from Riken Japan Health Science Foundation (3940), HeLa human cervical adenocarcinoma derived cells were obtained from ATCC (CCL-2, Manassas, Virginia), and AGS human gastric adenocarcinoma derived cells were obtained from ATCC (CRL-1739, Manassas, Virginia). AZ-521 and HeLa cells were maintained in MEM (Fisher Scientific), and AGS cells were maintained in DMEM (Fisher Scientific), each supplemented with 2 mM glutamine (Fisher Scientific), 10% FBS (Sigma-Aldrich), 100 mU/mL penicillin (Fisher Scientific), and 1 mg/mL streptomycin sulfate (Fisher Scientific). All cell lines were maintained at 37 °C, under 5% CO_2_ within a humidified atmosphere.

### Chemicals, reagents, and primary antibodies

Unless otherwise noted, all chemicals and reagents were purchased from Sigma-Aldrich. Primary antibodies against TOM20 (rabbit polyclonal, SC-11415), EEA1 (rabbit polyclonal, SC-33585), LAMP1 (mouse monoclonal, SC-20011), DRP1 (mouse monoclonal, BD-611112), and cytochrome *c* (mouse monoclonal, BD-556432) were obtained from BD Biosciences; and β-actin (rabbit monoclonal, CS-4970) were obtained from Cell Signaling Technology (Danvers, MA). The rabbit anti-VacA primary antibody was generated in lab(36).

### Purification of VacA

*Helicobacter pylori (Hp)* 60190 (*cag* PAI^+^, *vacA* s1/m1; ATCC 49503) was cultured in bisulfite/sulfite-free *Brucella* (BSFB) broth (4 mL, 0.5% NaCl, 0.2% β-cyclodextrin, 0.1% dextrose (all from Sigma-Aldrich), and, 1% peptone, 1% tryptone, and 0.2% yeast extract (all from BD Bacto), overlaid on 2% agar plates made of Ham’s F-12 medium (Sigma-Aldrich) supplemented with 5% fetal bovine serum (FBS) (Sigma-Aldrich)), and incubated in a microaerophilic environment (10% O_2_, 5% CO_2_).

After 48 h, the liquid culture phase was collected as a starter culture and diluted 1:100 in fresh BSFB (final volume of 2 L) supplemented with 5 µg/mL vancomycin (Sigma-Aldrich), and incubated on a shaker platform in a humidified, microaerophilic (10% O_2_, 5% CO_2_), 37 °C incubator. After 48 h, *Hp* cultures were centrifuged (5,000 x*g*, 4 °C, 30 min), the supernatants were collected, and VacA was precipitated by adding solid ammonium sulfate (Fisher Scientific) to 90% saturation (662 g/L) at 4 °C with constant stirring. After 4 h, the precipitated material was collected by centrifugation (5,000 x*g*, 4 °C, 30 min) and dissolved in “wash buffer” (10 mM Na_2_HPO_4_, pH 7.0; Sigma-Aldrich), and then dialyzed at 4 °C in the wash buffer (200 times the volume of the precipitate solution), with four buffer changes spanning 12 h, using 50 kDa molecular weight cutoff (MWCO) dialysis tubing (Spectra/Por 6 RC, Spectrum Chemical, New Brunswick, NJ). The dialyzed solution was cleared of any remaining insoluble material by centrifugation (5,000 x*g*, 4 °C, 30 min), and the supernatant was further cleared by filtration using a 0.22 μM MWCO filtration funnel (Stericup, EMD Millipore, Burlington, MA). The VacA-containing soluble fraction was loaded onto an anion exchange column (DEAE Sephacel, GE healthcare, Chicago, IL) pre-equilibrated with wash buffer. After washing the column with 3 bed volumes of wash buffer, VacA was eluted from the resin with wash buffer supplemented with 0.2 M NaCl and collected in 1 mL fractions. The purity of VacA within each fraction was assessed by SDS-PAGE and Coomassie Staining (G-250 Coomassie Brilliant Blue; Sigma-Aldrich). Fractions containing VacA without additional visible protein bands were combined and dialyzed for 12 h (with 4 buffer changes) at 4 °C in PBS (pH 7.4), using 50 kDa MWCO dialysis tubing. Following dialysis, purified VacA was filter-sterilized using a 0.22 μm pore syringe filter, aliquoted into single use vials, and stored at −20 °C. The total protein concentration was determined using the bicinchoninic acid (BCA) *assay* (Thermo Fisher Scientific).

### VacA exposure conditions

Monolayers of cells were incubated in the absence or presence of VacA (250 nM) under “cold” pulse conditions (4 ⁰C for 30 min), washed with PBS (pH 7.2), and incubated at 37 °C under 5% CO_2_ in fresh cell culture medium in the absence of VacA. Alternatively, AZ-521 cells were preloaded by incubating at 4 °C in the absence or presence of VacA (250 nM). After 30 min, cells were further incubated continuously at 37 °C. After 1, 4, or 24 h for both “cold” pulse and continuous exposure conditions, cells were analyzed for indicated phenotypes.

### Immunofluorescence microscopy

Following experimental treatments in chambered microscope slides, AZ-521, AGS, and HeLa cells were fixed in 5% formaldehyde at 37 °C for 15 min, washed with PBS (pH 7.4), permeabilized in 0.1% Triton-X 100 in PBS (pH 7.4) at room temperature for 15 min, washed with PBS pH 7.2, blocked with 5% bovine serum albumin (BSA; Sigma-Aldrich) for 1 h at room temperature, washed with 0.1% Tween-20 in PBS (pH 7.4), and probed for the mitochondrial marker TOM20 with an anti-TOM20 rabbit polyclonal antibody (1:1000 dilution; Sigma-Aldrich) and anti-cytochrome *c* mouse monoclonal antibody (1:500 dilution; BD Biosciences) in 1% BSA in PBS (pH 7.4) overnight at 4 °C, followed by Alexa Fluor-488 conjugated goat anti-rabbit secondary antibody (1:1000 dilution; Life Technologies, Carlsbad, CA), Alexa Fluor-488 conjugated goat anti-mouse secondary antibody (1:1000 dilution; Life Technologies), and/or Alexa Fluor-555 conjugated donkey anti-rabbit secondary antibody (1:1000 dilution; Life Technologies) in 1% BSA in PBS (pH 7.4) at room temperature for 2 h and counterstained with DAPI (0.5 µg/ml in PBS (pH 7.4), room temperature, 15 min), then mounted in ProLong Gold Antifade Reagent (Life Technologies) overnight and sealed. Cells were imaged by DIC epifluorescence microscopy using a DAPI and FITC filter on a Delta Vision RT microscope.

### Analysis of mitochondrial fragmentation

Mitochondrial filament lengths were measured manually using the Imaris imaging processing software (version 7.4.2). Mitochondria were analyzed from a single focal plane within 5 separate cells (∼63 mitochondria per cell, total of 8598 mitochondria analyzed). Data from three independent biological replicates each consisting of three technical replicates were combined and relativized to the untreated control.

### Analysis of mitochondrial transmembrane potential

AZ-521, AGS, and HeLa cells were incubated in tetramethylrhodamine ethyl ester perchlorate (TMRE) (10 nM, Life Technologies) during the final 30 min of each experiment. The cells were washed with PBS (pH 7.4), detached in 0.05% trypsin at 37 °C for 5 min, and collected in 10% FBS in PBS (pH 7.4) on ice. Relative TMRE fluorescence intensity was measured by flow cytometry using the FL2 channel (575/30-nm bandpass filter) on a BD FACSCanto II flow analyzer (BD Biosciences). Approximately 10,000 events were analyzed for each independent replicate. Data from three independent biological replicates each consisting of three technical replicates were combined and relativized to the untreated control.

### Analysis of ATP synthesis

AZ-521, AGS, or HeLa cells were cultured in glucose free, galactose (10 mM; Fisher) containing medium for 2 days prior to experimentation. ATP levels were measured using MitoTox Glo (Promega, Madison, WI), according to manufacturer’s instructions. The values were normalized by total protein, using the BCA assay (Thermo Fisher). Data from three independent biological replicates each consisting of three technical replicates were combined and relativized to the untreated control.

### Oxygen consumption measurements

AZ-521, AGS, or HeLa cells were cultured in a 96-well, black-walled cell culture plate (Corning Inc., Corning, NY) and adapted in glucose free, galactose-containing (10 mM; Fisher Scientific) medium for 2 days prior to experimentation. Following the completion of each experiment, cells were washed twice with PBS (pH 7.4), then incubated in serum-free media containing MitoXpress Xtra solution. Wells were overlaid with mineral oil and measurements were obtained at 380 nm for the excitation, and 650 nm for the emission at 37°C. The reciprocal values were normalized to total protein, using the BCA assay (Thermo Fisher). Data from three independent biological replicates each consisting of three technical replicates were combined and relativized to the untreated control.

### Lactic acid production

Extracellular lactic acid levels were used to determine the relative levels of aerobic glycolysis. AZ-521, AGS, or HeLa cells were cultured in a 96-well, black-walled cell culture plate (Corning Inc.) and adapted in glucose free, galactose-containing (10 mM; Fisher Scientific) medium for 2 days prior to experimentation. Following the completion of each experiment, culture supernatants were transferred to a fresh plate to perform the glycolysis assay using the Oxygen Consumption/Glycolysis Dual Assay Kit (Cayman Chemical, Ann Arbor, MI) according to the manufacturer’s instructions. Briefly, culture supernatants were combined with assay buffer, incubated at room temperature for 30 min, and analyzed for absorbance at 490 nm. The values were normalized to total protein, using the BCA assay (Thermo Fisher). Data from three independent biological replicates each consisting of three technical replicates were combined and relativized to the untreated control.

### Statistical analysis

All data were combined from three independent biological replicates, each comprising at least three technical replicates. For pairwise comparisons, significance was determined using a 2-tailed distribution, paired *t* test. For multiple comparisons, significance was determined using a one-way ANOVA with Tukey’s multiple comparisons test. Unless otherwise indicated, an alpha level of 0.05 was used as a threshold for statistical significance and all error bars represent standard error of the mean (SEM). Graphpad Prism (version 6.01) was used for all statistical analyses.

