## Supplemental Figure for "Restoration of mitochondrial structure and function within *Helicobacter pylori* VacA intoxicated cells"

**SUPPLEMENTAL FIGURES**

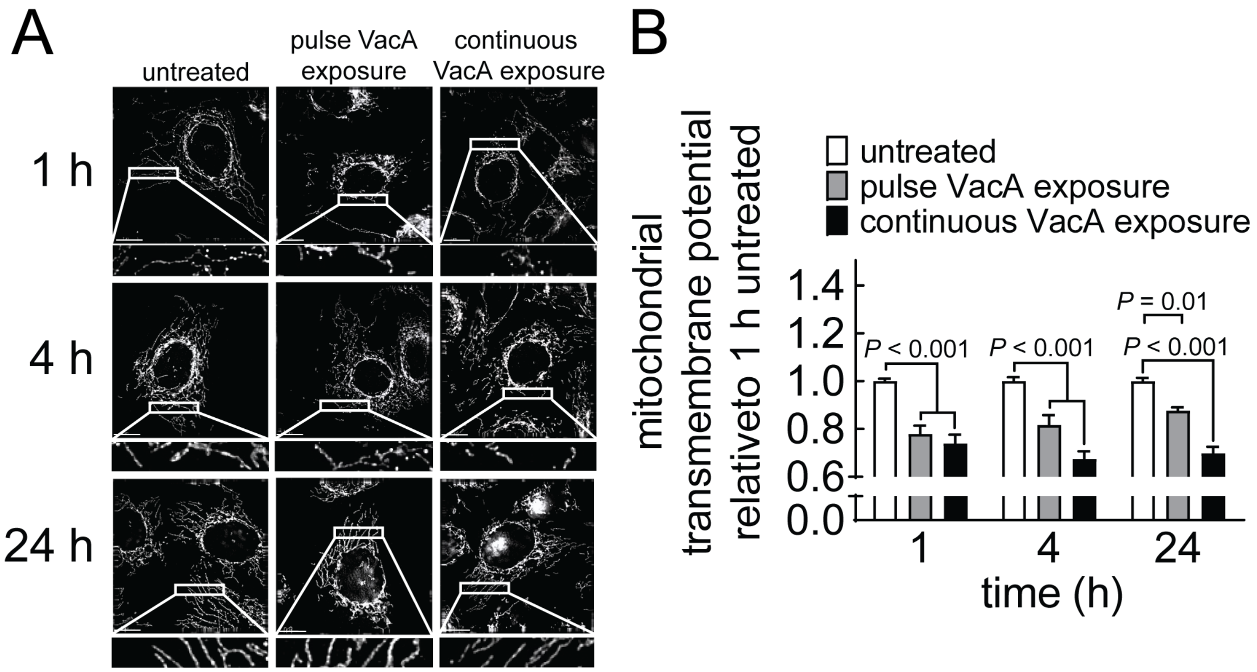


**Supplementary Figure 1. Mitochondrial structure recovery.** AGS cells were incubated in the absence or presence of VacA (250 nM) under “cold” pulse exposure conditions or continuous exposure conditions. After 1, 4, or 24 h, cells were collected for analyses. Mitochondria were stained with anti-TOM20 (A), followed by incubation with fluorescently labeled secondary antibodies, and imaged by fluorescence. Scale bars represent 10 µm. Results in (A) are representative of those collected in three independent biological replicates, each of which data are collected from a single focal plane within 5 separate cells. The mitochondrial transmembrane potential (B) was measured by flow cytometry with TMRE staining (10 nM, 30 min) and 10,000 events collected for each individual treatment. Data were combined from three independent experiments each performed in triplicate (B). Error bars represent standard error of the median. Statistical significance was determined by one-way ANOVA with an alpha of

0.001 ($\alpha$=0.001), using Tukey's correction for multiple comparisons against the untreated control within each timepoint.
